# *Plasmodium* infection disrupts the T follicular helper cell response to heterologous immunization

**DOI:** 10.1101/2022.09.20.508672

**Authors:** Mary F. Fontana, Erica Ollmann Saphire, Marion Pepper

## Abstract

Naturally acquired immunity to malaria develops only after many years and repeated exposures, raising the question of whether *Plasmodium* parasites, the etiological agents of malaria, suppress the ability of dendritic cells (DCs) to activate optimal T cell responses. We demonstrated recently that B cells, rather than DCs, are the principal activators of CD4+ T cells in murine malaria. In the present study, we further investigated factors that might prevent DCs from priming *Plasmodium*-specific T helper cell responses. We found that DCs were significantly less efficient at taking up infected red blood cells (iRBCs) compared to soluble antigen, whereas B cells more readily bound iRBCs. To assess whether DCs retained the capacity to present soluble antigen during malaria, we measured responses to a heterologous protein immunization administered to naïve mice or mice infected with *P. chabaudi*. Antigen uptake, DC activation, and expansion of immunogen-specific T cells were intact in infected mice, indicating DCs remained functional. However, polarization of the immunogen-specific response was dramatically altered, with a near-complete loss of germinal center T follicular helper cells specific for the immunogen, accompanied by significant reductions in antigen-specific B cells and antibody. Our results indicate that DCs remain competent to activate T cells during *Plasmodium* infection, but that T cell polarization and humoral responses are severely disrupted. This study provides mechanistic insight into the development of both *Plasmodium*-specific and heterologous adaptive responses in hosts with malaria.

## INTRODUCTION

Malaria, caused by parasites of the genus *Plasmodium*, is a leading global driver of infection-related mortality, resulting in 241 million cases of disease and over 600,000 deaths in 2020 (*1*). Clinical immunity to malaria (*i.e.* protection from symptoms) develops only after many years and repeated infections. Consequently, children in endemic areas remain vulnerable to severe illness and death for several years (*2–4*), making the slow and incomplete acquisition of immunity a matter of grave concern for human health. The mechanisms underlying this apparently poor development of immunity have been the subject of intense research for decades. Antigenic diversity among *Plasmodium* parasites is likely one factor; additionally, however, evidence suggests that immune responses generated in hosts with malaria may be defective or sub-optimal, relative to responses against immunizations or other pathogens (*5*). For example, antibodies against *P. falciparum* in naturally exposed children are short-lived relative to antibodies against unrelated vaccine antigens (*5, 6*), and *P. falciparum* infection has been linked to diminished T helper cell responses in children and adults (*7–11*). These observations raise the question of whether *Plasmodium* parasites subvert optimal immune responses to facilitate repeated infection of their mammalian hosts. The immune defects associated with *Plasmodium* infection are not limited to responses against the parasite itself: children with acute malaria exhibit impaired responses to a range of vaccinations (*12, 13*), and *Plasmodium* infection causes declines in pre-existing, unrelated vaccine-induced antibodies and plasma cells in both mice and humans (*14, 15*). From a public health perspective, it is therefore also important to understand how malaria negatively impacts heterologous immunizations.

One proposed mechanism for the development of sub-optimal or short-lived immune responses in malaria is that *Plasmodium* parasites may suppress the activity of dendritic cells (DCs), innate immune cells that initiate canonical T cell responses to microbes (evidence for malaria-induced dysfunction reviewed in (*16*)). Some studies have found that *P. falciparum*-infected RBCs (iRBCs) or parasite products can render human DCs refractory to activation and restrict priming of CD4+ T cells *in vitro* (*17–22*), and humans with malaria have fewer and less activated circulating DCs than healthy controls (*23, 24*). However, these decreases in circulating DCs during infection could reflect recruitment to infected tissues, and additional *in vitro* studies have concluded that DCs *do* become activated and can prime T cells after exposure to iRBCs and parasite products (*25–27*). Overall, the question of whether malaria suppresses DC activity remains a subject for debate (*16, 28*).

Mouse models have provided additional characterizations of DC function during *Plasmodium* infection, but with similarly conflicting results. Some studies have reported that *Plasmodium* suppresses DC function under specific circumstances (*29–31*), but others have established that DCs isolated from infected mice can present parasite-derived antigens to activate T cells and generate protective responses (*18*, *29*, *30*, *32*–*34*). Whether they actually perform this function in hosts with malaria is less clear. Several papers have sought to test a role for DCs *in vivo* using a transgenic mouse model that permits deletion of CD11c-expressing cells. These studies concluded that conventional CD11c^hi^ DCs were important for antigen presentation and T cell responses in malaria (*35, 36*). However, CD11c is also upregulated on activated B cells, making the results of these studies difficult to interpret. Using more selective genetic tools to ablate antigen presentation specifically in DCs or B cells, we recently demonstrated that B cells, rather than DCs, are the dominant antigen-presenting cells (APCs) that direct activation and polarization of the CD4+ T cell response in mice with malaria (*37*). In light of this finding, we considered it important to revisit the question of DC dysfunction in mice with malaria.

Historically, the ability of B cells to present antigen to T cells has been studied primarily in the context of cognate interactions with CD4+ T cells that have already been activated by DCs. However, B cells can also interact with and activate naïve T cells directly. Hong et al. showed recently that whereas DCs were required for activating CD4+ T cell responses to a soluble protein antigen, T cell responses to the same antigen delivered on a nanoparticle required presentation by B cells (*38*). In light of this finding, here we examined whether a differential ability to take up particle-like iRBCs might bias presentation of *Plasmodium* parasite antigens away from DCs and towards B cells. Further, we tested whether DCs in *Plasmodium*-infected mice remained competent to activate T cell responses to a heterologous, soluble protein antigen–an approach that allowed us to assess the functional capacity of DCs in hosts with malaria, independent of whether DCs can efficiently obtain iRBC-associated antigen in vivo. We found that DCs are relatively poor phagocytes of iRBCs, which may be one contributing factor in the dominance of B cells as APCs in this disease. We also found that DCs remained competent to activate CD4+ T cell responses to soluble antigens in hosts with malaria, suggesting no global dysfunction occurs. However, CD4+ T cell polarization, germinal center formation, and antibody production were profoundly altered, a finding that has implications for vaccination schemes in malaria-endemic regions.

## RESULTS

### DCs do not efficiently capture infected RBCs in vivo

We recently demonstrated that mice infected with the rodent-adapted parasite *P. yoelii* developed an antigen-specific CD4+ T cell response that was strongly polarized to a T follicular helper cell (Tfh) phenotype, specialized for assisting B cells with affinity maturation within germinal centers. In mice with malaria, these antigen-specific T cells possessed diminished proliferative capacity and longevity compared to Th cells specific for the same epitope delivered in the context of an LCMV infection. Using genetically modified mice to selectively disrupt or restore antigen presentation in either DCs or B cells, we also determined that B cells, rather than DCs, were the primary APCs in this infection and orchestrated the striking Tfh polarization phenotype (*37*).

We wished to better understand factors that favor B cells as the dominant APCs in this infection model, especially in light of recent work demonstrating that different APCs are required for soluble versus nanoparticle-associated antigen (*38*). Since *Plasmodium* infects RBCs, its antigens are primarily associated with “particles” *in vivo* (i.e. RBCs) rather than existing in soluble form. We hypothesized that the particulate nature of the parasite antigen itself might bias the APC response toward B cells and away from DCs, perhaps contributing to DCs’ apparent lack of participation in antigen presentation to T cells in this infection (*37*). A previous study found that DCs do interact with iRBCs in the spleens of infected mice (*35*), and indeed, it is well-established that DCs from infected mice *can* activate *Plasmodium*-specific T cell responses *ex vivo*, indicating that they must take up some antigen from iRBCs (*18*, *30*, *32*–*34*). Yet the fact that they are capable of T cell activation *ex vivo* does not necessarily mean that they are the preferred or dominant APCs activating T cells *in vivo*, as infection could impact DC localization or migration. Some of these studies also examined B cells *ex vivo* and concluded that they did not activate CD4+ T cells during malaria; however, they specifically isolated the CD11c- subset, likely excluding the activated B cells that are most likely to serve as APCs (*33, 34*). Other studies either did not specifically examine B cells or deliberately excluded them from analysis. Given our recent findings on the central role of B cells in activating CD4+ T cell responses in this system, we considered it critical to revisit previous work examining uptake of iRBCs in the spleens of infected mice.

Modifying the experimental approach of Borges da Silva et al. (*35*), we used flow cytometry to examine how well various splenic APC subsets, including B cells, could bind to labeled RBCs infected with the parasite *P. chabaudi*. Like *P. falciparum*, *P. chabaudi* exports membrane proteins that render infected RBCs (iRBCs) cytoadherent (*39*), a characteristic found to be important for interaction with (and suppression of) DCs in one study (*17*). We enriched iRBCs from the blood of infected mice (Fig. S1A), labeled them with a fluorescent dye, and injected them intravenously into naïve mice. Differentially labeled naïve RBCs (nRBCs) were injected for comparison. We harvested spleens after 30 minutes and assessed association of labeled nRBCs and iRBCs with various splenic subsets, including B cells (defined as CD45+ Thy1.2- MHCII+ B220+ CD11c^lo/int^), DCs (CD45+ Thy1.2- MHCII^hi^ CD11c^hi^), and red pulp macrophages (RPMs; CD45+ Thy1.2- MHCII+ F4/80^hi^), which specialize in uptake and recycling of senescent RBCs (Fig. 1A). We found that DCs made up only a small fraction of total RBC+ splenocytes; instead, RBCs were mostly distributed between RPMs and B cells (Fig. 1B–D). Additional flow experiments examining Siglec-H+ plasmacytoid DCs did not detect significant association of iRBCs with these cells (data not shown). The frequencies of iRBC+ DCs that we measured were similar to those reported by Borges da Silva et al. (*35*); however, that group did not perform a comparison with B cells.

**Figure 1.**
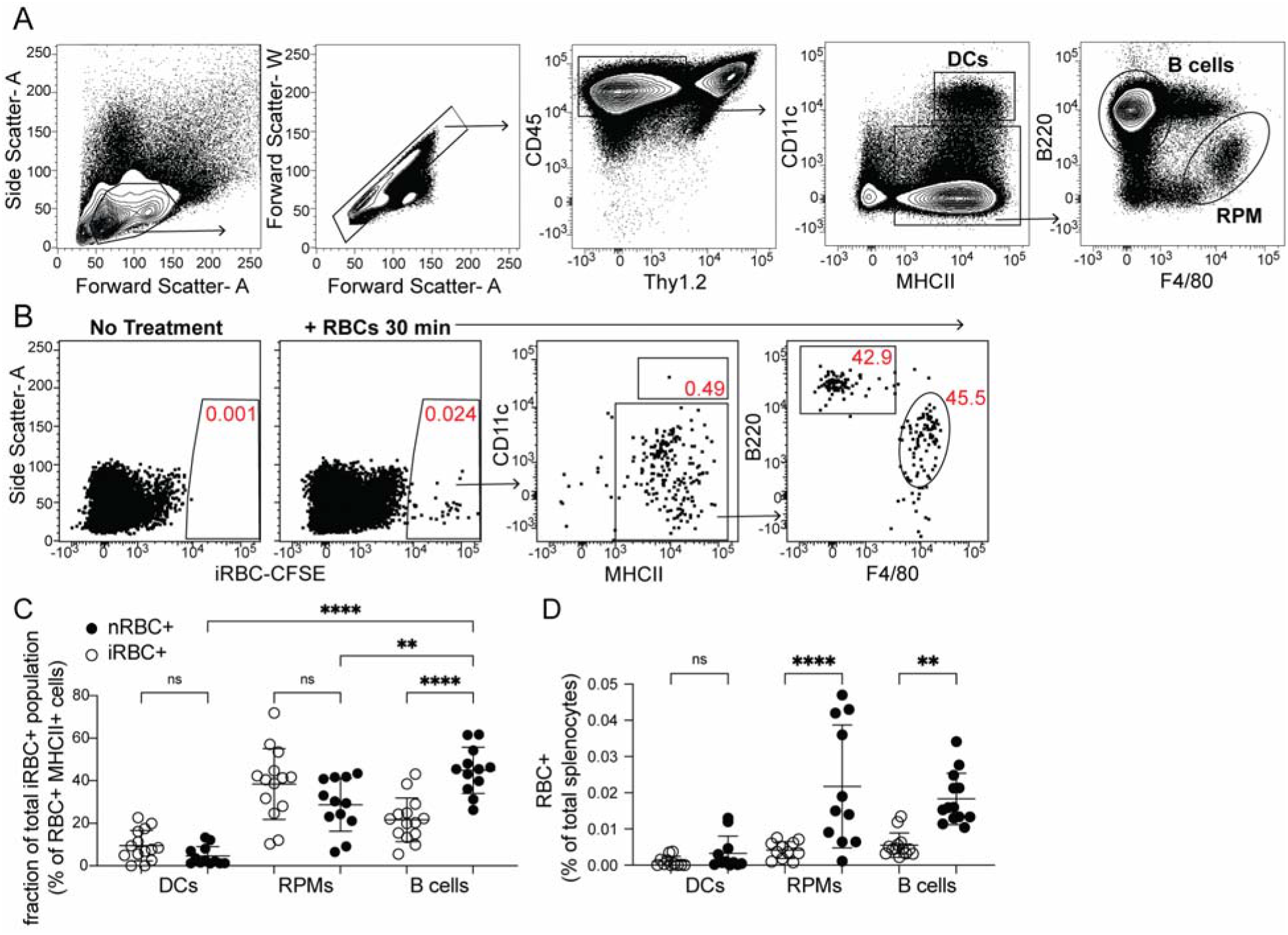
DCs capture only a small fraction of infected RBCs in vivo. Red blood cells infected with *Plasmodium chabaudi* (iRBCs) were enriched from infected mice, labeled with a fluorescent dye, and injected intravenously into naïve C57Bl/6J mice. Labeled naïve RBCs (nRBCs) were injected as a control. Splenocytes were analyzed 30 min later for fluoresecent iRBC label by flow cytometry. (A) Representative gating of bulk dendritic cells (DCs), B cells, and red pulp macrophages (RPMs). (B) Representative plots showing the distribution of labeled iRBCs in DCs, B cells, and RPMs. (C) Quantification showing the percentage of the total RBC+ population that consisted of DCs, RPMs, or B cells as indicated. (D) Quantification of RBC+ DCs, RPMs, and B cells expressed as a percentage of total splenocytes. A and B depict representative plots from one of four independent experiments. C and D show pooled data (mean +/− SD) from all 4 experiments, with each symbol representing the indicated subset from one mouse (n = 12 mice for iRBC treatment, 14 for nRBC treatment). **, p < 0.01 and ****, p < 0.0001 by one-way ANOVA with Tukey’s post-test. n.s., not significant.

In our *in vivo* experiments, DCs bound similar frequencies of nRBCs and iRBCs (Fig. 1D). Interestingly, B cells bound significantly more iRBCs than nRBCs both *in vivo* and *in vitro* (Figs. 1C–D, S1B), although some apparent RBC binding to B cells *in vivo* was artifactual and occurred during processing (Fig. S1C, D). This preferential binding of iRBCs was independent of the B cell receptor (Fig. S1B), leading us to speculate that B cells may selectively interact with iRBCs through other, antigen-independent pathways such as complement receptors, Fc receptors, or scavenger receptors (*40, 41*). Altogether, these experiments show that of all the iRBCs interacting with APCs in the spleen, only a small percentage are associated with DCs.

### DCs take up soluble antigen more efficiently than infected RBCs

Because DCs are well-established as the primary APCs that activate naïve T cells in response to soluble antigens, we next tested how DC uptake of iRBCs compared to uptake of soluble protein. We simultaneously injected the easily traceable fluorescent molecule phycoerythrin (PE) along with labeled iRBCs intravenously into mice and measured uptake by splenic APC subsets after 30 minutes. Gating separately on all MHCII^+^ cells that carried either the PE or the iRBC label, we further divided the labeled cells into APC populations (Fig. 2A). A majority of both PE^+^ and iRBC^+^ events were B cells, reflecting the abundance of this subset, which comprises ~50% of splenocytes. However, DCs accounted for around 10% of PE+ events, despite comprising only 1-2% of splenocytes. In contrast, the iRBC^+^ population included very few DCs (1-3%, commensurate with their total splenic frequency), but many more RPMs (~20%) in addition to B cells (Fig.2A, B).

**Figure 2.**
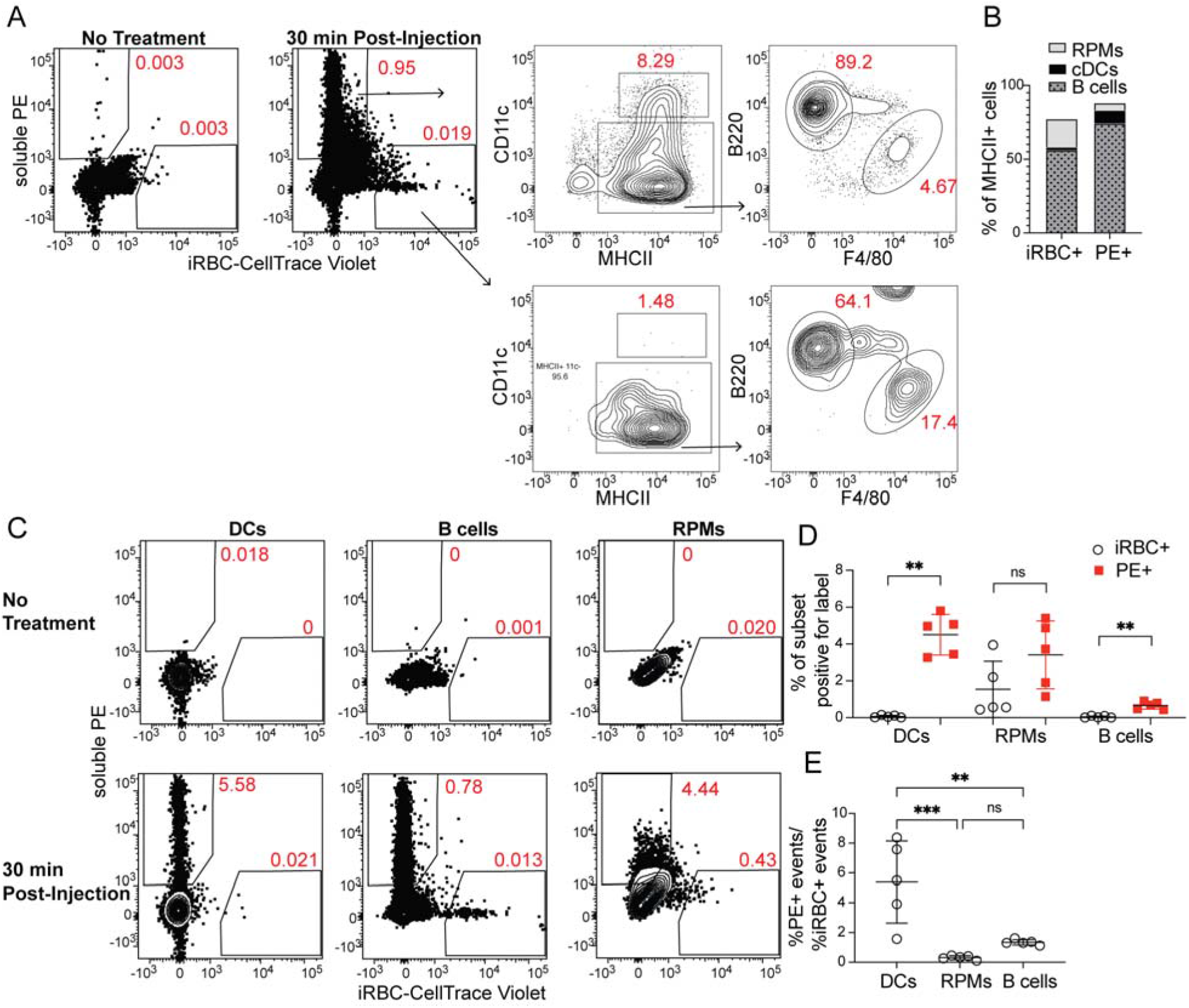
DCs take up soluble antigen more efficiently than infected RBCs. Mice were intravenously injected with enriched, fluorescently labeled iRBCs together with PE and splenocytes were analyzed 30 min later for acquisition of fluorescent signal. (A) Representative gating of PE+ or iRBC+ splenocytes, further subsetted into the indicated APC populations. (B) Quantification of the distribution of PE and iRBC labeling among DCs, B cells, and RPMs. (C) Alternative gating strategy showing DCs, B cells, and RPMs in untreated mice (top row) and 30 min after injection of PE and iRBCs (bottom panel). (D) Quantification showing the percentage of each APC subset that acquired iRBC or PE label, as gated in (C). (E) Ratio of the percentage of each subset that acquired PE signal to the percentage of that subset that acquired iRBC signal. B, D, and E show pooled data from two independent experiments (n = 5). **, p < 0.01 and ***, p < 0.001 by one-way ANOVA with Tukey’s post-test. n.s., not significant.

As a second way of analyzing these same data, we examined the fraction of each APC subset that acquired either PE or iRBC label. Whereas ~5% of CD11c^hi^ MHCII^hi^ splenic DCs had taken up IV-injected PE, the percentage of DCs labeling with iRBCs was virtually zero (Fig. 2C, D). We considered that this difference in uptake could be due to different effective doses of PE and iRBCs. However, DCs capture a greater fraction of the total cell-associated PE than the total cell-associated iRBC signal, indicating a better ability to capture PE (Fig. 2A, B). Further, the ratio of PE+ DCs to iRBC+ DCs within each mouse was significantly higher than the ratio of PE+ RPMs or B cells to iRBC+ RPMs or B cells, respectively, indicating that DCs are biased toward taking up PE more so than iRBCs, relative to the other APC subsets examined (Fig. 2E). Thus, DCs do not capture iRBCs as efficiently as soluble antigen *in vivo*. Instead, the majority of iRBCs associate with other APC subsets, including B cells. It is possible that even a small amount of antigen uptake and presentation is sufficient to permit activation of naïve T cells, and indeed it has been shown that DCs from infected mice do acquire *Plasmodium* antigens *in vivo* (*18*, *30*, *32*–*34*). However, in light of our previous study identifying B cells as the primary APCs in mice with malaria (*37*), as well as data showing that B cells preferentially present nanoparticle-associated antigens to T cells (*38*), we suggest that poor acquisition of iRBC-borne antigen may be one factor underlying the lack of substantial contribution from DCs to the antigen-specific CD4+ T cell response in *Plasmodium*-infected mice.

### Monocyte-derived DCs take up majority of soluble antigen in Plasmodium-infected mice

Our initial experiments raised the possibility that differential uptake of parasite antigen might contribute to the prominence of B cells in antigen presentation during malaria. However, they did not address the question of whether DC activation, antigen presentation, upregulation of costimulatory molecules, and/or secretion of cytokines and chemokines are suppressed during *Plasmodium* infection, as has been suggested (*16*, *17*, *27*, *30*). To probe the question of whether DCs retain the capacity to present antigen and activate T cells in mice with malaria, independent of their ability to access or take up particulate antigen, we investigated antigen uptake, DC activation state, and initiation of T cell responses to a heterologous, soluble protein antigen delivered to naïve or *P. chabaudi*-infected mice.

First, we tested whether DC uptake of a soluble protein was altered in infected mice. We injected PE into naïve mice or mice infected for 5 days with *P. chabaudi*. At this timepoint, *Plasmodium*-specific T cell differentiation is under way (*37*) but germinal centers, which are delayed in malaria relative to their kinetics following protein immunization, have not yet formed (*42*). As expected, we found that parasitemia was still ascending (Fig. 3A) but total DC numbers in the spleen were not significantly different from naïve mice (Fig. 3B). 30 minutes after injection, we observed equivalent or slightly higher frequencies of PE+ DCs in infected compared to uninfected mice (Fig. 3C), indicating that there is no defect in acquisition of soluble antigen by DCs during infection. We next examined whether infection altered the phenotype or identity of PE+ DCs, since DC populations change during infection. Specifically, we measured the frequency of newly recruited monocyte-derived DCs (moDCs), which were defined as MHCII^hi^ CD11c^hi^ CD11b^hi^ CD64+ (Fig. 3D). moDCs were rare in the spleens of uninfected mice. In infected mice, they comprised ~30% of all DCs, but made up ~80% of PE+ DCs (Fig. 3D, E). Thus, moDCs disproportionately capture the majority of soluble antigen in infected mice, compared to CD64- conventional DCs (cDCs).

**Figure 3.**
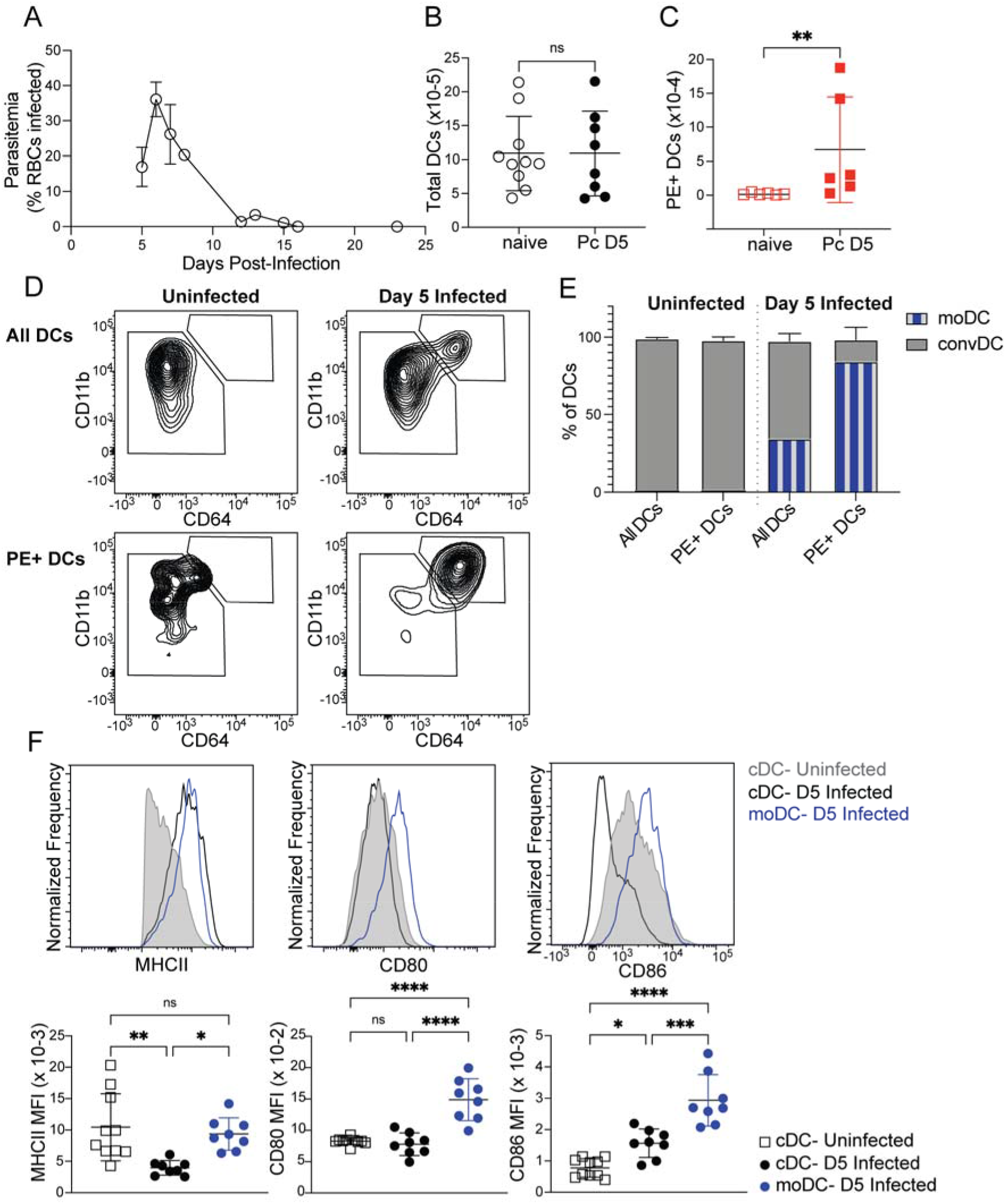
Monocyte-derived DCs take up majority of soluble antigen in *Plasmodium*-infected mice. (A) Mice were infected with 10^6^ *P. chabaudi*-parasitized RBCs and parasitemia was monitored by thin blood smear. (B) MHC^hi^ CD11c^hi^ DCs were enumerated by flow cytometry in the spleens of mice at homeostasis or 5 days after *P. chabaudi* infection (Pc D5). (C) Naïve or day 5-infected mice were injected intraperitoneally (i.p.) with PE, and PE+ splenic DCs were quantified after 30 min by flow cytometry. (D) Representative gating of conventional CD64- DCs (cDC) and CD11b^hi^ CD64+ monocyte-derived DCs (moDC) in naïve or infected mice injected with PE. Top plots show all DCs, while bottom plots show PE+ DCs. (E) Plots depicting the percentage of total or PE+ DCs that are cDC or moDC in naïve or infected mice. (F) Histograms and quantification showing expression (median fluorescent intensity, MFI) of MHCII, CD80, and CD86 on cDC and moDC. moDC are shown only from infected mice due to their scarcity in naïve mice. A shows data from one experiment (n = 3), representative of many. B, C, E and F show mean +/− SD from at least three experiments (n = 6 per group for C; 10 uninfected, 8 infected for all others). Red numbers next to flow plot gates represent the frequency of cells within the gate. *, p < 0.05, **, p < 0.01, ***, p < 0.001, ****, p < 0.0001 by Mann-Whitney (B, C) or one-way ANOVA with Tukey’s post-test (all others). n.s., not significant.

Given this observation, we next questioned whether moDCs possessed the components to activate T cells. Flow cytometric analysis of surface markers revealed that moDCs expressed at least as much MHCII as cDCs, and had significantly higher expression of the costimulatory molecules CD80 and CD86 compared to cDCs from either uninfected or infected mice (Fig. 3F). Altogether, our data demonstrate that DC uptake of soluble antigen remains intact after 5 days of *P. chabaudi* infection, but that the antigen-capturing subset shifts primarily from cDCs to moDCs. These moDCs express robust levels of MHCII and costimulatory molecules, and therefore possess the basic requirements for efficient T cell priming.

### Intact CD4+ T cell expansion, but disrupted polarization, following heterologous immunization in infected mice

We next examined whether the DCs that took up soluble antigens in infected mice retained the capacity to activate CD4+ T cells *in vivo*. For these experiments, we used a recombinant form of the glycoprotein (GP) from lymphocytic choriomeningitis virus (LCMV) because it contains the well-defined CD4+ T cell epitope GP66, permitting CD4+ T cells specific for this epitope to be detected with a fluorescently labeled peptide-MHCII tetramer (*43*). We previously characterized GP66-specific CD4+ T cell responses in mice infected with a transgenic *P. yoelii* parasite that expresses this epitope (*37*); thus, we have extensive data on the phenotype and kinetics of GP66-specific CD4+ T cells when the epitope is delivered in the context of an iRBC. In the present study, administration of GP with adjuvant during an infection with (non-transgenic) *P. chabaudi* allowed us to examine the same population of GP66-specific T cells as they responded to a soluble antigen presented by DCs, all within the context of a malaria-inflamed spleen.

We injected uninfected or day 5-infected mice with GP plus adjuvant and analyzed splenic GP66-specific CD4+ T cells by flow cytometry 8 days after immunization (which was 13 days post-infection). GP66-specific T cells were rare in unimmunized mice but expanded dramatically following immunization in both uninfected and infected mice. No T cell expansion was observed in infected mice that did not receive GP. The total number of GP66-specific CD4+ T cells was not significantly different between uninfected and infected mice following immunization of both, indicating that the ability of DCs to present soluble antigen and activate T cells was intact in infected mice (Fig. 4A, B).

**Figure 4.**
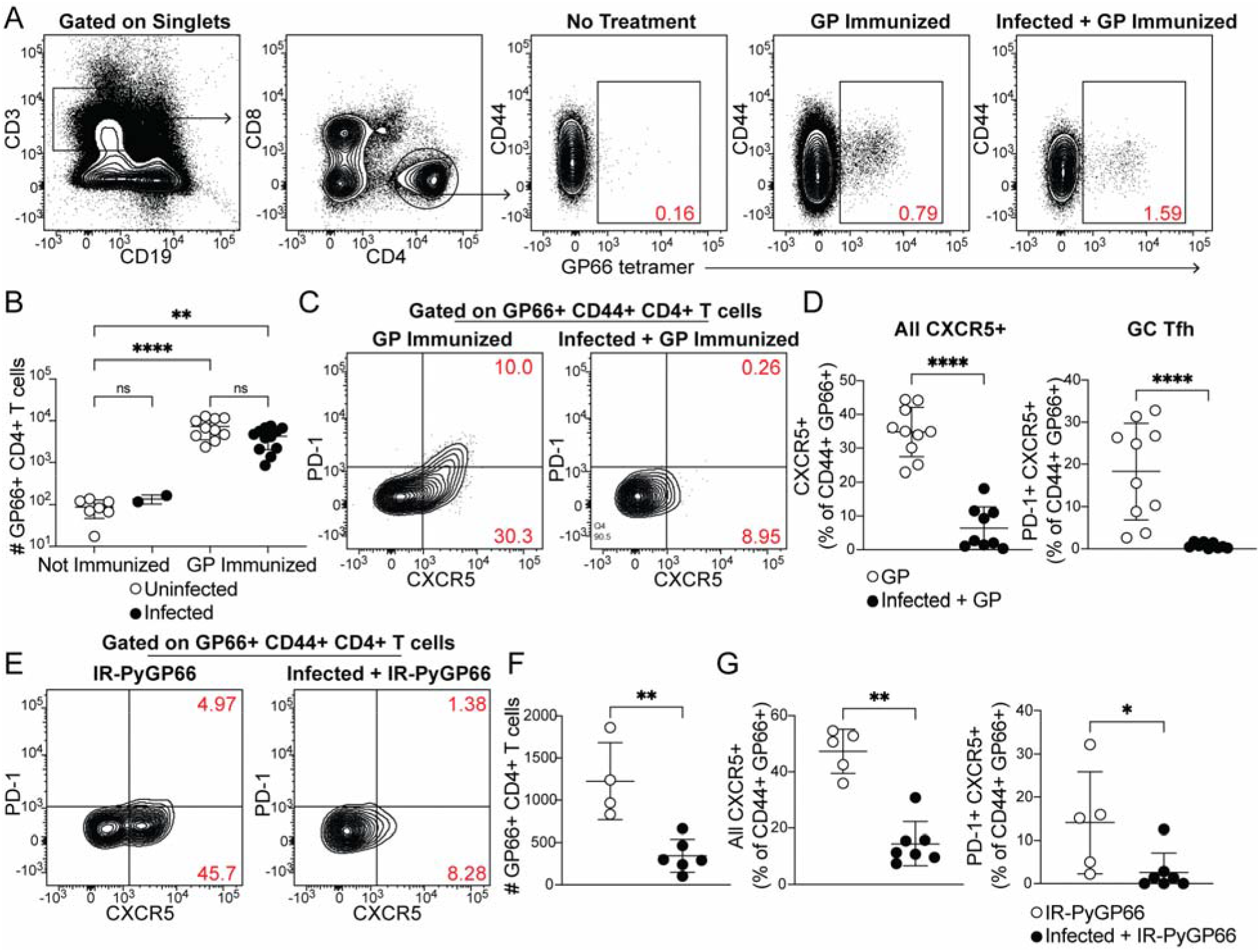
CD4+ T cell expansion, but disrupted polarization, following heterologous immunization in infected mice. (A-D) Mice were immunized i.p. with recombinant LCMV glycoprotein (GP) plus adjuvant, either alone (GP Immunized) or 5 days after *P. chabaudi* infection (Infected + GP Immunized). GP66-specific CD4+ T cells were analyzed in spleen 8 days after immunization. (A) Representative gating and (B) quantification of GP66+ CD4+ T cells from uninfected and infected mice with or without GP immunization. (C) Gating and (D) quantification of activated Tfh (CD44^hi^ CXCR5+) and GC-Tfh (CD44^hi^ CXCR5+ PD-1^hi^) GP66-specific cells. (E-G) Mice left uninfected or infected for 5 days with *P. chabaudi* were immunized with irradiated *P. yoelii*-parasitized RBCs expressing GP66 (IR-PyGP66). (E) Representative flow plot of CXCR5 and PD-1 expression on GP66-specific T cells 8 days after IR-PyGP66 immunization. (F-G) Quantification of total GP66-specific CD4+ T cells (F) and Tfh and GC Tfh frequencies (G) 8 days after immunization. A-D represent data from four independent experiments (pooled n = 10 “GP” mice, 12 “Infected + GP” mice). F-G show pooled results from two independent experiments (n = 5 immunized, 7 infected + immunized). Red numbers within flow plot gates represent the frequency of cells within the gate. *, p < 0.05, **, p < 0.01 and ****, p < 0.0001 by one-way ANOVA with Tukey’s post-test (B) or Mann-Whitney (others). n.s., not significant.

We next examined the phenotype of GP66-specific T cells in uninfected and infected mice. Specifically, we measured polarization of CD4+ T cells into CXCR5+ PD-1^hi^ germinal center (GC) Tfh cells. As expected, uninfected mice developed a distinct population of GC Tfh cells, as well as a sizeable subset of CXCR5+ PD-1^lo^ Tfh cells. Strikingly, however, the Tfh and GC Tfh populations were virtually absent among GP66-specific cells in infected mice following GP immunization (Fig. 4C, D). This was not due to a generalized loss of Tfh cells in infected mice, since abundant CXCR5+ T cells were observed in the bulk (non-tetramer-binding) T cell population (Fig. S2) and our previous study found strong Tfh polarization of GP66-specific CD4^+^ T cells at similar timepoints after infection with GP66+ parasites (*37*). Together, these data demonstrate that an established *P. chabaudi* infection does not prevent expansion of T cells in response to a heterologous soluble antigen, but does abrogate the polarization of antigen-specific T cells into B cell-helping Tfh and GC Tfh cells.

Previous work has shown that DCs can orchestrate initial differentiation of CXCR5+ Tfh cells 2-4 days after immunization, but that maintenance of the Tfh phenotype at day 8 requires both cognate and non-cognate interactions with B cells (*44–46*). Therefore we hypothesized that interactions between DCs and T cells were intact in infected mice, allowing for T cell expansion, but that infection disrupted interactions between GP-specific T cells and B cells, preventing establishment of Tfh and GC Tfh populations. To further dissect B-T cognate interactions in the infected spleen, we treated uninfected mice, or mice infected for 5 days with *P. chabaudi*, with irradiated RBCs containing nonreplicating transgenic GP66+ *P. yoelii* parasites (*47*) and measured GP66-specific T cell responses 8 days later. Since iRBC-associated GP66 is primarily presented by B cells (*37*), we reasoned that if interactions between newly activated B and T cells were disrupted in day 5-infected mice, expansion of GP66-specific CD4+ T cells should be defective. This was indeed what we observed (Fig. 4E, F). CXCR5+ Tfh and GC Tfh subsets were also significantly diminished in infected mice, consistent with the loss of cognate and non-cognate B-T interactions that normally induce Tfh polarization (Fig. 4E, G). Taken together, our data suggest that in an infected spleen, DCs remain competent for uptake and presentation of soluble antigens, but that the signals required for T cell polarization into Tfh cells–including both cognate and non-cognate interactions with B cells–are disrupted.

### B cell expansion and GC formation in response to heterologous immunization are curtailed in infected mice

Because antigen-specific GC Tfh cell responses are sharply diminished when antigen is administered during an established *P. chabaudi* infection (Fig. 4C, D), we examined expansion of antigen-specific B cells and development of the GC B cell population after the same immunization scheme. A fluorescently labeled GP protein tetramer was used to identify GP-specific B cells (Fig. 5A) (*48*). These cells expanded robustly in number 8 days after GP immunization of uninfected mice, but not following GP immunization of mice infected for 5 days with *P. chabaudi* (Fig. 5A, B). The frequency of CD138+ GP-specific plasmablasts was modest 8 days post-immunization and was not different between infected and uninfected mice (Fig. 5C). In contrast, a sizeable population of GP-specific GC B cells (CD138- CD38− GL7+) appeared in uninfected, immunized mice, but was greatly decreased in infected mice following immunization (Fig. 5D). Previous work from our group found that *Plasmodium*-specific GC B cells are readily detected at this time (i.e., 13 days after infection) (*42*). These data identify a functional consequence of the drastic decrease in GP66-specific GC Tfh cells in infected mice, as Tfh cells are required to sustain the repeated rounds of proliferation and selection that antigen-specific B cells undergo in GCs. Consistent with the decreases in total and GC B cell numbers, GP-specific serum IgG antibody titers were much lower in infected compared to uninfected mice 7 days after immunization (Fig. 5E and F). A slight but significant defect in serum IgG persisted 14 days after immunization; however, after 3 weeks, titers in infected mice increased to the levels of those in uninfected mice. This timepoint corresponds roughly to the time that *P. chabaudi* parasitemia is cleared to sub-patent levels, although a low-grade chronic infection persists for several months in this model (*39*). These data show that established *Plasmodium* infection significantly decreases expansion of B cells, establishment of germinal centers, and production of antibodies specific for a heterologous protein antigen.

**Figure 5.**
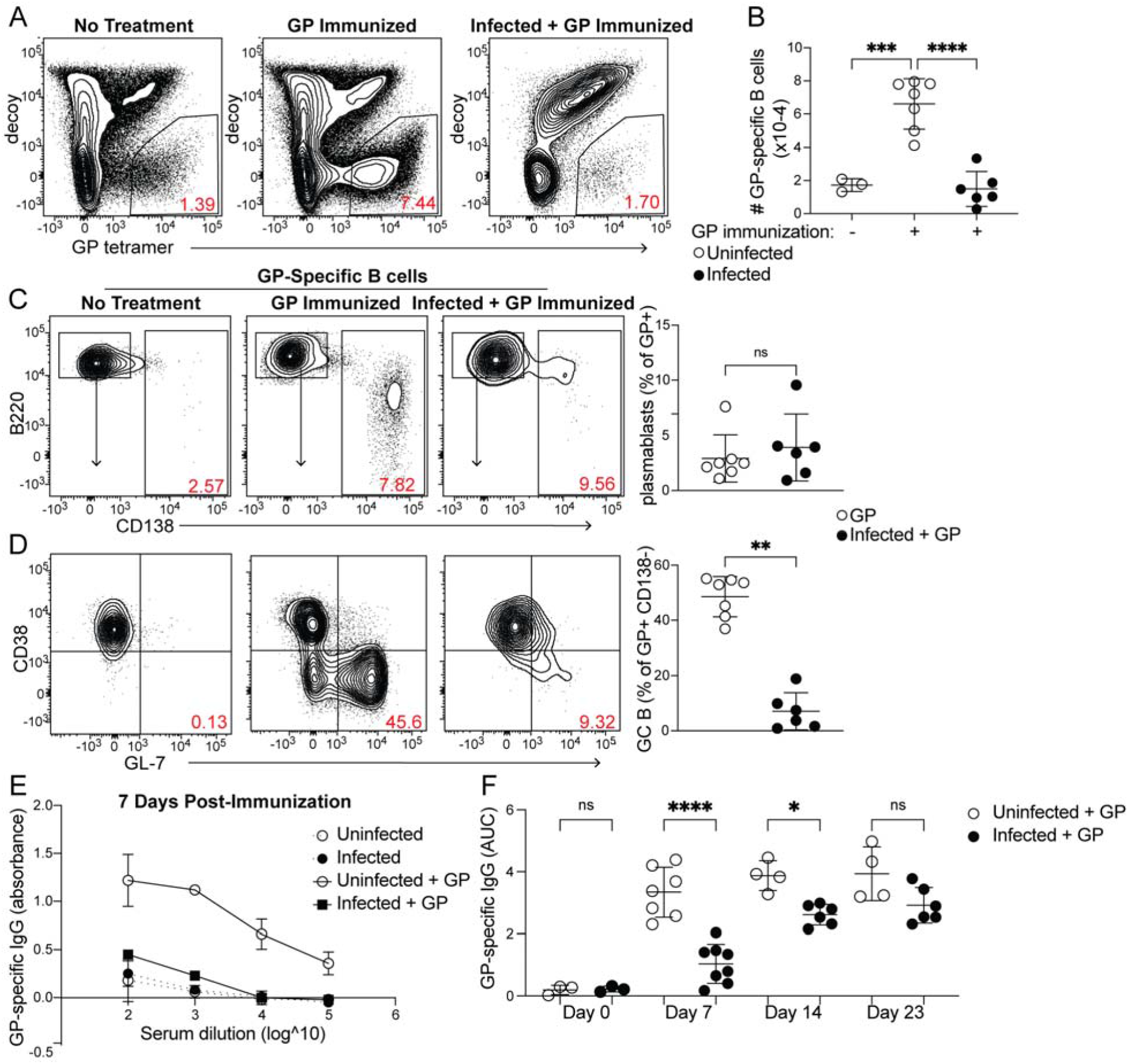
Curtailed B cell expansion and antibody production in infected mice following heterologous immunization. Mice were immunized with or without concurrent *P. chabaudi* infection, as in Fig. 4. (A) Representative gating and (B) quantification of GP-specific splenic B cells 8 days post-immunization. In A, plots were pre-gated on Thy1.2- B cells. (C, D) Gating (left panels) and quantification (right) of CD138+ plasmablasts (C) or CD138- CD38− GL7+ GC B cells (D) 8 days after immunization with GP. (E) GP-specific IgG antibody was measured by ELISA in the serum of uninfected or infected mice without immunization or 7 days after immunization with GP. (F) GP-specific serum antibodies (expressed as Area Under the Curve, AUC) measured by ELISA at the indicated times post-immunization. A-C represent data from three independent experiments (n = 7 GP, 6 Infected + GP); E and F are pooled from two experiments. Red numbers within flow plot gates represent the frequency of cells within the gate. *, p < 0.05, **, p < 0.01, ***, p. 0.001, and ****, p < 0.0001 by one-way ANOVA with Tukey’s post-test (B, F) or Mann-Whitney (others). n.s., not significant.

## DISCUSSION

Our results paint a nuanced picture of DC function during *Plasmodium* infection. DCs do not take up iRBCs efficiently in vivo, compared to their uptake of soluble antigen. This relatively poor ability to capture RBC-associated antigen may contribute to the importance of B cells as APCs in this infection setting, since the majority of iRBCs were found to associate with B cells, along with RPMs. However, differential uptake is likely not the only factor biasing antigen presentation toward B cells, since DCs do acquire sufficient antigen in infected mice to activate CD4+ T cells ex vivo (*18*, *30*, *32*–*34*).

Although they did not capture iRBCs efficiently, DCs in both uninfected and infected mice readily took up soluble antigen and expressed the Class II and costimulatory molecules necessary to activate CD4+ T cells. Further, although the composition of the DC subset changed during infection, the magnitude of the T cell response to soluble protein immunization was similar in both uninfected and infected mice, indicating that DCs were competent to activate CD4+ T cells in a *Plasmodium*-infected spleen. The use of different markers to define DC subsets complicates direct comparisons between our findings and previous studies, but Sponaas et al. identified CD8− CD11b+ DCs from infected mice as the most robust activators of CD4+ T cells *ex vivo* (*32*); we believe that at the timepoints examined, this subset largely overlaps with the CD11b+ CD64+ moDC population that we identified here as the main DC subset capturing soluble antigen in infected mice. In other contexts, moDCs have been found fully capable of antigen presentation and T cell activation (*49, 50*).

Altogether, our findings are consistent with previous studies showing that DCs isolated from infected mice are capable of activating CD4+ T cells *in vitro*. In addition, though, our work provides a readout of interactions between DCs and CD4+ T cells within malarial hosts rather than *ex vivo*. This is important because the splenic architecture undergoes profound alteration in mice and humans with malaria, with dissolution of marginal zones and blurring of borders between the T cell zones and B cell follicles; it has been suggested that these architectural disruptions contribute to the observed delay in establishment of germinal centers, and subsequent failure to make optimal antibody responses (*51–56*). These tissue changes have already begun to occur 5 days after infection, when we immunized mice with GP, and reach their peak during the window in which the GP-specific response is being initiated (*51*, *52*, *55*, *57*). Our data suggest that even within this dramatically altered tissue environment, DCs are able to interact with both antigen and CD4+ T cells in a way that permits robust expansion of the latter, ruling out the existence of global DC dysfunction in the malaria-inflamed spleen.

At the same time, we did observe a near-complete defect in differentiation of GP-specific Tfh and GC Tfh cells in mice that were immunized with GP 5 days after *Plasmodium* infection. We and others previously showed that ICOS-mediated interactions between B and T cells support maintenance of Tfh cells, while cognate B-T interactions are necessary for the GC Tfh subset (*45*). In light of this, we infer that the loss of GP-specific Tfh and GC Tfh cells in *Plasmodium*-infected mice is due to disruption of both cognate and non-cognate interactions between GP-specific T cells and B cells. Here, alteration of the splenic architecture may play an important role, with the blurring of the T-B border potentially making it more difficult for activated T cells to locate B cells. The inflammatory milieu of the infected spleen may also negatively impact Tfh development; we and others have found that blocking inflammatory cytokines enhances the differentiation of Tfh cells and GC B cells during infections with other strains of Plasmodium (*56, 57*). Finally, it is interesting to speculate whether the BCR-independent binding of iRBCs that we observed may dilute or misdirect the antigen-specific B cell response in hosts with malaria. Indeed, *Plasmodium* infection is known to induce polyclonal, nonspecific B cell activation as well as activation of B cells autoreactive for RBC antigens (*58–61*). This could also contribute to the failure of GP-specific B cells to expand and support the differentiation of cognate Tfh cells.

The initial motivation for this study was to investigate mechanisms underlying our previous discovery that B cells, rather than DCs, served as the dominant APC population shaping the CD4+ T cell response during *Plasmodium* infection. By expanding previously established assays of iRBC uptake to include examination of B cells, we found that the majority of iRBCs associate with splenic B cells rather than with DCs, and that B cells preferentially bind iRBCs over nRBCs. This finding is consistent with a previous study showing that B cells were the primary APCs that activated CD4+ T cell responses when antigen was delivered in nanoparticle form, whereas DCs were key for initiating responses to the same antigen in soluble form (*38*). In that study, B cell uptake of nanoparticle-associated antigen was BCR-dependent and performed by antigen-specific B cells. Here, we found that B cells can preferentially bind iRBCs independent of the BCR, although this does not preclude additional BCR-dependent binding by B cells specific for parasite antigens, such as those we have characterized extensively in previous studies (*42*, *48*, *56*, *62*). This selective but BCR-independent binding is intriguing; it suggests the existence of antigen-nonspecific interactions between iRBCs and B cells that might explain the reported activation of nonspecific polyclonal or autoreactive B cells during malaria. The literature suggests a number of possible such interaction pathways: receptors expressed by B cells might bind antibody, complement, exposed phosphatidylserines or parasite proteins on the iRBC surface (*40*, *63*, *64*). Once taken up, iRBCs contain immunostimulatory ligands, such as parasite RNA and DNA, that could activate B cells through Toll-like receptor signaling, alone or in conjunction with signaling from any of the above pathways (*65–68*). Further dissection of interactions between iRBCs and B cells therefore remains an important avenue for future research.

One important aspect of our study with relevance for human health is its insight into the detrimental effects of ongoing *Plasmodium* infection on antibody responses to immunization. Such effects have been appreciated for decades, most notably through measurement of poor responses to childhood vaccinations in children from endemic regions (*12–14*). Malaria also may affect antibody avidity (*55*). But the mechanisms underlying these malaria-associated decreases in antibody titers have mostly remained unknown, although one paper did identify increased turnover of serum antibodies as a contributing factor (*61*). We recently found that blood-stage-induced inflammation disrupts the development of sporozoite-specific B cells and antibodies in mice (*56*), perhaps providing an explanation for the relative scarcity of CSP-specific memory B cells in naturally exposed humans (*69, 70*). In the present study, we have further elucidated the cellular basis for defective antibody production during malaria: we showed that immunization during acute *Plasmodium* infection activates immunogen-specific CD4+ T cells, but fails to elicit the Tfh cells necessary to support a robust immunogen-specific B cell response, correlating with diminished B cell expansion and decreased serum antibody titers. The detrimental effect on antibody production is transient, suggesting that interactions between B and T cells eventually recover as the infection is cleared to sub-patent levels. However, in endemic areas, many clinically immune humans carry patent levels of parasites in their blood during much of the year (*4*). It remains to be seen whether asymptomatic parasitemia affects vaccine efficacy similarly to acute malaria. Even so, a strategy in which vaccine is administered together with anti-*Plasmodium* chemoprophylaxis has recently shown great promise (*71*), raising our hope that a greater understanding of malaria’s effects on the immune system will enable continued improvement of human protection from this and other diseases.

---

**Figure S1.**
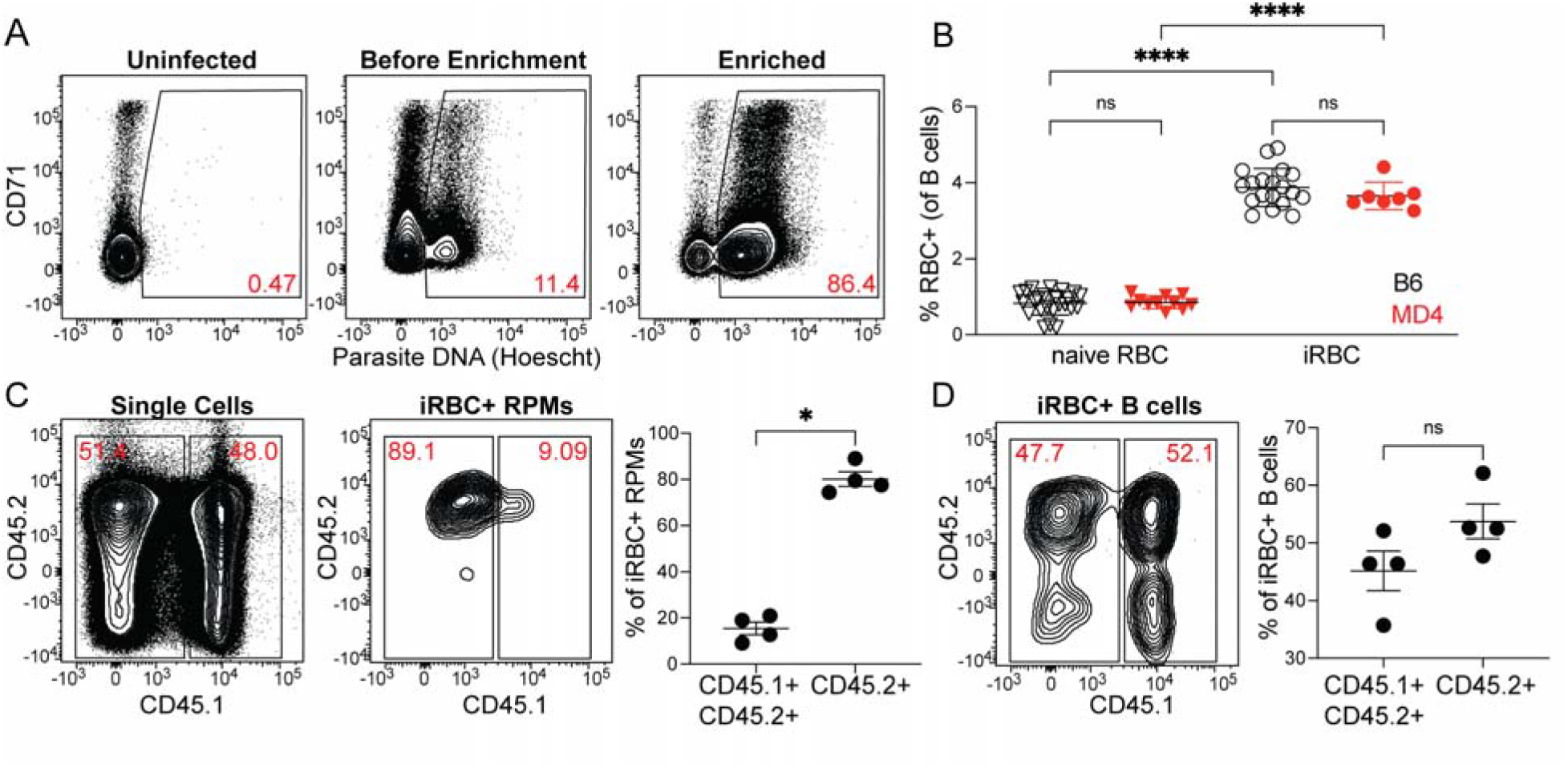
Uptake of enriched iRBCs by B cells. (A) Infected RBCs were enriched over a magnetic column and degree of enrichment was assessed by flow cytometry. An uninfected sample is shown for comparison to pre-enrichment and post-samples from an infected mouse. Plots shown are pre-gated on Ter119+ CD45+ cells; then iRBCs are identified by staining of nucleic acids with Hoescht. CD71 is used to identify reticulocytes. (B) B6 or MD4 splenocytes were incubated with labeled naive RBCs or enriched iRBCs and RBC binding to B cells was assessed by flow cytometry. (C-D) Labeled iRBCs were injected into CD45.2+ mice. Each spleen was harvested after 30 min and disrupted along with a spleen from an untreated CD45.1+ CD45.2+ mouse. Binding of iRBCs to splenic CD45+ cells was assessed by flow cytometry. (C) Contribution of each donor spleen to the total splenocyte pool (left panel) and subset of iRBC+ RPMs (middle panel). iRBC+ RPMs from each donor spleen are quantified (right panel). (D) Flow plot (left) and quantification (right) of the proportion of iRBC+ B cells derived from each donor spleen. A depicts représentative plots from one of many enrichments. B-D show représentative plots and pooled quantification from two independent experiments. *, p < 0.05 and ****, p < 0.0001 by one-way ANOVA with Tukey’s post-test (B) or Mann-Whitney (C, D). n.s., not significant.

**Figure S2.**
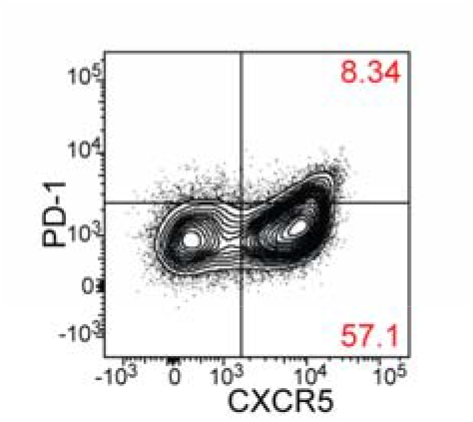
Abundant CXCR5 expression by bulk CD4+ T cells in infected mice. Représentative flow plot showing CXCR5 and PD-1 expression on bulk CD19- CD44hi CD4+ T cells in mice infected for 13 days with P. chabaudi. This timepoint corresponds to the time of analysis for GP66-specific T cells in mice immunized with GP 5 days after P. chabaudi infection, then harvested 8 days after immunization (Figure 4).

## METHODS

### Mice

Wild-type C57Bl/6J (B6), CD45.1+, and MD4 mice were purchased from Jackson Laboratories. B6 and CD45.1+ mice were crossed in-house to generate CD45.1+ CD45.2+ mice. All mice were group-housed at the University of Washington on a twelve-hour light-dark cycle under specific pathogen-free conditions. 8-12 week old male and female mice were used for all experiments. Mice were age- and sex-matched within each experiment, and no significant differences were observed between sexes. Mice were randomly assigned to experimental groups. Sample sizes were determined by past experience. No mice were excluded from analysis. All mouse experiments were conducted with the approval of the UW Institutional Animal Care and Use Committee in accordance with the guidelines of the NIH Office of Laboratory Animal Welfare.

### Plasmodium *parasites*

Wild-type *Plasmodium chabaudi* AS was obtained from the MR4 Stock Center and passaged in B6 mice. Transgenic *P. yoelii* 17XNL-GP66 has been described (*47*). For *P. chabaudi* infections, mice were injected intraperitoneally (i.p.) with 106 infected RBCs. Parasitemia was monitored by thin blood smear stained with Giemsa (*72*). For immunization with irradiated parasites, *P. yoelii* GP66-infected RBCs were suspended in Alsevers solution and subjected to 30,000 rad on ice. 107 irradiated iRBC were then injected i.p. into mice.

### Enrichment and labeling of iRBCs

Blood was collected by cardiac puncture from infected mice 6-7 days post-infection, when parasitemia was ~20-40%, and immediately diluted into 10mL PBS. It was then spun, resuspended in PBS, and loaded onto pre-washed LD columns (Miltenyi). The bound fraction was washed with 4-5mL PBS, eluted, spun down and resuspended at 5*107 RBC/mL in PBS. CFSE or CellTrace Violet (Invitrogen) was added to a final concentration of 5uM and cells were labeled for 20 min at room temperature. Labeled RBCs were washed in five volumes of cold DMEM + 10% FBS, spun, and resuspended in PBS at 10^8^ cells/mL for injection. Degree of enrichment was measured both by thin blood smear and by flow cytometry using Hoescht nucleic acid dye to identify iRBCs. To generate naïve RBCs, blood from a naïve mouse was collected and passed through an LD column. The flowthrough was collected and labeled as described for iRBCs. Injections of 10^8^ labeled RBCs were performed via retroorbital vein. In some cases, labeled iRBCs were mixed with 30ug PE (Agilent) prior to injection.

### Detection of RBC binding during processing of spleens

CD45.2+ (B6) mice were injected with labeled, enriched iRBCs as described above. After 30 min, each spleen was removed and placed into 5mL PBS on ice along with a freshly excised spleen from an untreated CD45.1+ CD45.2+ mouse. After harvests were complete, each pair of spleens was manually disrupted together, filtered through nylon mesh, and placed back on ice. Flow cytometric analysis was performed as described below.

### In vitro RBC binding

Spleens were aseptically removed from naïve B6 and MD4 mice after euthanasia according to approved methods. Single-cell suspensions were made, and 2*10^6^ B6 or MD4 splenocytes were mixed with 10^7^ labeled, enriched iRBCs or nRBCs in DMEM containing 10% FBS. The combined cells were incubated for 20 min at 37 degrees, followed by ACK lysis of RBCs and flow cytometric analysis (see below).

### Protein immunizations

10ug recombinant LCMV glycoprotein in PBS was diluted 1:1 in Sigma Adjuvant System (Sigma) and administered i.p. B and T cell responses were analyzed in spleens 8 days after immunization.

### Flow cytometry

For experiments involving analysis of DCs and macrophages, excised spleens were cut into 6-8 pieces and digested for 15 min at 37 degrees in digest buffer (RPMI + 2% FBS + 10mM HEPES + 1mg/mL Collagenase IV + 20ug/mL DNase I). The tissue was then manually disrupted and passed through nylon mesh to obtain single-cell suspensions. The digest step was omitted when only lymphocytes were analyzed. RBCs were lysed in ACK buffer and samples were blocked with anti-CD16/CD32 prior to labeling with antibodies. To analyze antigen-specific T cells, splenocytes were labeled with I-A(b) LCMV GP 66-77 tetramer (NIH Tetramer Core Facility) conjugated to APC (*37*). To analyze GP-specific B cells, splenocytes were incubated first with a decoy reagent labeled in-house with APC-Dylight755 (*73*), then with a fluorescently labeled GP tetramer (*48*). Magnetic anti-APC beads (Miltenyi) were used to enrich tetramer-positive cells after labeling (*42, 73*). Following enrichment, cells were incubated with antibodies and washed. Data were collected on an LSRII or Symphony (BD) and analyzed using FlowJo software (Treestar). Antibodies are listed in the Key Resources Table.

### ELISA

Blood was collected from the submental veins of mice and serum was snap-frozen until analysis. High-binding 96 well plates (Costar) were coated with recombinant GP (5ug/mL in PBS). After washing, serially diluted serum was applied to plates and incubated at room temperature for two hours or overnight at 4°C. Plates were washed and biotinylated anti-mouse IgG (1:1000; Biolegend) was added for 1h at room temperature, followed by streptavidin-HRP (1:500; Jackson) for 30 min at room temperature. After a final wash, plates were developed with TMB (Invitrogen), the reaction was stopped with 1M HCl, and absorbance was measured at 450nm. AUC was calculated with Graphpad Prism 9 software.

### Statistical Analysis

All statistics were calculated using Prism 9 software. The statistical tests used are listed in each figure legend.

## Data Availability

All data generated and analyzed in this study are contained within the paper, figures, and supporting files.

## ACKNOWLEDGEMENTS

We thank Kathryn Hastie for help with GP production and members of the Pepper lab for helpful discussions.

## AUTHOR CONTRIBUTIONS

MFF conceptualized the study, designed and performed experiments, analyzed data, and wrote the paper. EOS contributed GP protein. MP conceptualized the study, provided funding and supervision, and edited the paper.

## REFERENCES

1. World Malaria Report 2021, (available at https://www.who.int/teams/global-malaria-programme/reports/world-malaria-report-2021).

2. R. E. Black, S. Cousens, H. L. Johnson, J. E. Lawn, I. Rudan, D. G. Bassani, P. Jha, H. Campbell, C. F. Walker, R. Cibulskis, T. Eisele, L. Liu, C. Mathers, Child Health Epidemiology Reference Group of WHO and UNICEF, Global, regional, and national causes of child mortality in 2008: a systematic analysis. Lancet. 375, 1969–1987 (2010).

3. L. Schofield, I. Mueller, Clinical immunity to malaria. Curr. Mol. Med. 6, 205–221 (2006).

4. V. Ryg-Cornejo, A. Ly, D. S. Hansen, Immunological processes underlying the slow acquisition of humoral immunity to malaria. Parasitology. 143, 199–207 (2016).

5. S. Portugal, S. K. Pierce, P. D. Crompton, Young lives lost as B cells falter: what we are learning about antibody responses in malaria. J. Immunol. 190, 3039–3046 (2013).

6. P. D. Crompton, M. A. Kayala, B. Traore, K. Kayentao, A. Ongoiba, G. E. Weiss, D. M. Molina, C. R. Burk, M. Waisberg, A. Jasinskas, X. Tan, S. Doumbo, D. Doumtabe, Y. Kone, D. L. Narum, X. Liang, O. K. Doumbo, L. H. Miller, D. L. Doolan, P. Baldi, P. L. Felgner, S. K. Pierce, A prospective analysis of the Ab response to Plasmodium falciparum before and after a malaria season by protein microarray. Proc. Natl. Acad. Sci. U.S.A. 107, 6958–6963 (2010).

7. M. Ho, H. K. Webster, S. Looareesuwan, W. Supanaranond, R. E. Phillips, P. Chanthavanich, D. A. Warrell, Antigen-specific immunosuppression in human malaria due to Plasmodium falciparum. J. Infect. Dis. 153, 763–771 (1986).

8. M. Ho, H. K. Webster, B. Green, S. Looareesuwan, S. Kongchareon, N. J. White, Defective production of and response to IL-2 in acute human falciparum malaria. J. Immunol. 141, 2755–2759 (1988).

9. C. Chizzolini, G. Grau, A. Geinoz, D. Schrijvers, T Lymphocyte Interferon-Gamma Production Induced by Plasmodium Falciparum Antigen Is High in Recently Infected Non-Immune and Low in Immune Subjects. Clinical and experimental immunology. 79(1990),, doi:10.1111/j.1365-2249.1990.tb05133.x.

10. M. Rhee, B. Akanmori, M. Waterfall, E. Riley, Changes in Cytokine Production Associated With Acquired Immunity to Plasmodium Falciparum Malaria. Clinical and experimental immunology. 126(2001),, doi:10.1046/j.1365-2249.2001.01681.x.

11. P. Bejon, G. Warimwe, C. L. Mackintosh, M. J. Mackinnon, S. M. Kinyanjui, J. N. Musyoki, P. C. Bull, K. Marsh, Analysis of Immunity to Febrile Malaria in Children That Distinguishes Immunity from Lack of Exposure. Infect. Immun. 77, 1917–1923 (2009).

12. B. M. Greenwood, A. M. Bradley-Moore, A. D. Bryceson, A. Palit, Immunosuppression in children with malaria. Lancet. 1, 169–172 (1972).

13. W. A. Williamson, B. M. Greenwood, Impairment of the immune response to vaccination after acute malaria. Lancet. 1, 1328–1329 (1978).

14. S. Banga, J. D. Coursen, S. Portugal, T. M. Tran, L. Hancox, A. Ongoiba, B. Traore, O. K. Doumbo, C.-Y. Huang, J. T. Harty, P. D. Crompton, Impact of acute malaria on pre-existing antibodies to viral and vaccine antigens in mice and humans. PLoS One. 10, e0125090 (2015).

15. D. H. L. Ng, J. J. Skehel, G. Kassiotis, J. Langhorne, Recovery of an antiviral antibody response following attrition caused by unrelated infection. PLoS Pathog. 10, e1003843 (2014).

16. M. N. Wykes, M. F. Good, What really happens to dendritic cells during malaria? Nat Rev Microbiol. 6, 864–870 (2008).

17. B. C. Urban, D. J. Ferguson, A. Pain, N. Willcox, M. Plebanski, J. M. Austyn, D. J. Roberts, Plasmodium falciparum-infected erythrocytes modulate the maturation of dendritic cells. Nature. 400, 73–77 (1999).

18. D. S. Pouniotis, O. Proudfoot, V. Bogdanoska, V. Apostolopoulos, T. Fifis, M. Plebanski, Dendritic Cells Induce Immunity and Long-Lasting Protection against Blood-Stage Malaria despite an In Vitro Parasite-Induced Maturation Defect. Infection and Immunity. 72, 5331–5339 (2004).

19. S. R. Elliott, T. P. Spurck, J. M. Dodin, A. G. Maier, T. S. Voss, F. Yosaatmadja, P. D. Payne, G. I. McFadden, A. F. Cowman, S. J. Rogerson, L. Schofield, G. V. Brown, Inhibition of dendritic cell maturation by malaria is dose dependent and does not require Plasmodium falciparum erythrocyte membrane protein 1. Infect. Immun. 75, 3621–3632 (2007).

20. X. Z. Yap, R. J. Lundie, G. Feng, J. Pooley, J. G. Beeson, M. O’Keeffe, Different Life Cycle Stages of Plasmodium falciparum Induce Contrasting Responses in Dendritic Cells. Front Immunol. 10, 32 (2019).

21. O. R. Millington, C. Di Lorenzo, R. S. Phillips, P. Garside, J. M. Brewer, Suppression of adaptive immunity to heterologous antigens during Plasmodium infection through hemozoin-induced failure of dendritic cell function. J. Biol. 5, 5 (2006).

22. A. D. Pack, P. V. Schwartzhoff, Z. R. Zacharias, D. Fernandez-Ruiz, W. R. Heath, P. Gurung, K. L. Legge, C. J. Janse, N. S. Butler, Hemozoin-mediated inflammasome activation limits long-lived anti-malarial immunity. Cell Rep. 36, 109586 (2021).

23. A. Pinzon-Charry, T. Woodberry, V. Kienzle, V. McPhun, G. Minigo, D. A. Lampah, E. Kenangalem, C. Engwerda, J. A. López, N. M. Anstey, M. F. Good, Apoptosis and dysfunction of blood dendritic cells in patients with falciparum and vivax malaria. J. Exp. Med. 210, 1635–1646 (2013).

24. T. Woodberry, G. Minigo, K. A. Piera, F. H. Amante, A. Pinzon-Charry, M. F. Good, J. A. Lopez, C. R. Engwerda, J. S. McCarthy, N. M. Anstey, Low-level Plasmodium falciparum blood-stage infection causes dendritic cell apoptosis and dysfunction in healthy volunteers. J. Infect. Dis. 206, 333–340 (2012).

25. C. Coban, K. J. Ishii, T. Kawai, H. Hemmi, S. Sato, S. Uematsu, M. Yamamoto, O. Takeuchi, S. Itagaki, N. Kumar, T. Horii, S. Akira, Toll-like receptor 9 mediates innate immune activation by the malaria pigment hemozoin. J. Exp. Med. 201, 19–25 (2005).

26. J. W. Griffith, T. Sun, M. T. McIntosh, R. Bucala, Pure Hemozoin is inflammatory in vivo and activates the NALP3 inflammasome via release of uric acid. J. Immunol. 183, 5208–5220 (2009).

27. A. Götz, M. S. Tang, M. C. Ty, C. Arama, A. Ongoiba, D. Doumtabe, B. Traore, P. D. Crompton, P. Loke, A. Rodriguez, Atypical activation of dendritic cells by Plasmodium falciparum. Proc. Natl. Acad. Sci. U.S.A. 114, E10568–E10577 (2017).

28. S. S. Struik, E. M. Riley, Does malaria suffer from lack of memory? Immunol Rev. 201, 268–290 (2004).

29. M. N. Wykes, X. Q. Liu, L. Beattie, D. I. Stanisic, K. J. Stacey, M. J. Smyth, R. Thomas, M. F. Good, Plasmodium strain determines dendritic cell function essential for survival from malaria. PLoS Pathog. 3, e96 (2007).

30. A.-M. Sponaas, N. Belyaev, M. Falck-Hansen, A. Potocnik, J. Langhorne, Transient deficiency of dendritic cells results in lack of a merozoite surface protein 1-specific CD4 T cell response during peak Plasmodium chabaudi blood-stage infection. Infect Immun. 80, 4248–4256 (2012).

31. M. N. Wykes, X. Q. Liu, S. Jiang, C. Hirunpetcharat, M. F. Good, Systemic tumor necrosis factor generated during lethal Plasmodium infections impairs dendritic cell function. J Immunol. 179, 3982–3987 (2007).

32. A.-M. Sponaas, E. T. Cadman, C. Voisine, V. Harrison, A. Boonstra, A. O’Garra, J. Langhorne, Malaria infection changes the ability of splenic dendritic cell populations to stimulate antigen-specific T cells. J Exp Med. 203, 1427–1433 (2006).

33. C. Voisine, B. Mastelic, A.-M. Sponaas, J. Langhorne, Classical CD11c+ dendritic cells, not plasmacytoid dendritic cells, induce T cell responses to Plasmodium chabaudi malaria. Int. J. Parasitol. 40, 711–719 (2010).

34. J. A. Perry, A. Rush, R. J. Wilson, C. S. Olver, A. C. Avery, Dendritic cells from malaria-infected mice are fully functional APC. J. Immunol. 172, 475–482 (2004).

35. H. Borges da Silva, R. Fonseca, A. D. A. Cassado, É. Machado de Salles, M. N. de Menezes, J. Langhorne, K. R. Perez, I. M. Cuccovia, B. Ryffel, V. M. Barreto, C. R. F. Marinho, S. B. Boscardin, J. M. Álvarez, M. R. D’Império-Lima, C. E. Tadokoro, In vivo approaches reveal a key role for DCs in CD4+ T cell activation and parasite clearance during the acute phase of experimental blood-stage malaria. PLoS Pathog. 11, e1004598 (2015).

36. K. Ueffing, H. Abberger, A. M. Westendorf, K. Matuschewski, J. Buer, W. Hansen, Conventional CD11chigh Dendritic Cells Are Important for T Cell Priming during the Initial Phase of Plasmodium yoelii Infection, but Are Dispensable at Later Time Points. Front Immunol. 8, 1333 (2017).

37. E. N. Arroyo, M. Pepper, B cells are sufficient to prime the dominant CD4+ Tfh response to Plasmodium infection. J Exp Med. 217, e20190849 (2020).

38. S. Hong, Z. Zhang, H. Liu, M. Tian, X. Zhu, Z. Zhang, W. Wang, X. Zhou, F. Zhang, Q. Ge, B. Zhu, H. Tang, Z. Hua, B. Hou, B Cells Are the Dominant Antigen-Presenting Cells that Activate Naive CD4+ T Cells upon Immunization with a Virus-Derived Nanoparticle Antigen. Immunity. 49, 695–708.e4 (2018).

39. R. Stephens, R. L. Culleton, T. J. Lamb, The contribution of Plasmodium chabaudi to our understanding of malaria. Trends Parasitol. 28, 73–82 (2012).

40. R. Vijay, J. J. Guthmiller, A. J. Sturtz, S. Crooks, J. T. Johnson, L. Li, L. Y.-L. Lan, R. L. Pope, Y. Chen, K. J. Rogers, N. Dutta, J. E. Toombs, M. E. Wilson, P. C. Wilson, W. Maury, R. A. Brekken, N. S. Butler, Hemolysis-associated phosphatidylserine exposure promotes polyclonal plasmablast differentiation. J Exp Med. 218, e20202359 (2021).

41. L. Shen, S. Tenzer, W. Storck, D. Hobernik, V. K. Raker, K. Fischer, S. Decker, A. Dzionek, S. Krauthäuser, M. Diken, A. Nikolaev, J. Maxeiner, P. Schuster, C. Kappel, A. Verschoor, H. Schild, S. Grabbe, M. Bros, Protein corona-mediated targeting of nanocarriers to B cells allows redirection of allergic immune responses. J Allergy Clin Immunol. 142, 1558–1570 (2018).

42. A. T. Krishnamurty, C. D. Thouvenel, S. Portugal, G. J. Keitany, K. S. Kim, A. Holder, P. D. Crompton, D. J. Rawlings, M. Pepper, Somatically Hypermutated Plasmodium-Specific IgM(+) Memory B Cells Are Rapid, Plastic, Early Responders upon Malaria Rechallenge. Immunity. 45, 402–414 (2016).

43. R. A. Zander, R. Vijay, A. D. Pack, J. J. Guthmiller, A. C. Graham, S. E. Lindner, A. M. Vaughan, S. H. I. Kappe, N. S. Butler, Th1-like Plasmodium-Specific Memory CD4+ T Cells Support Humoral Immunity. Cell Reports. 21, 1839–1852 (2017).

44. S. M. Kerfoot, G. Yaari, J. R. Patel, K. L. Johnson, D. G. Gonzalez, S. H. Kleinstein, A. M. Haberman, Germinal center B cell and T follicular helper cell development initiates in the interfollicular zone. Immunity. 34, 947–960 (2011).

45. M. Pepper, A. Pagán, B. Igyártó, J. Taylor, M. Jenkins, Opposing Signals From the Bcl6 Transcription Factor and the interleukin-2 Receptor Generate T Helper 1 Central and Effector Memory Cells. Immunity. 35(2011),, doi:10.1016/j.immuni.2011.09.009.

46. T. von der Weid, J. Langhorne, Altered response of CD4+ T cell subsets to Plasmodium chabaudi chabaudi in B cell-deficient mice. Int Immunol. 5, 1343–1348 (1993).

47. W. O. Hahn, N. S. Butler, S. E. Lindner, H. M. Akilesh, D. N. Sather, S. H. Kappe, J. A. Hamerman, M. Gale, W. C. Liles, M. Pepper, cGAS-mediated control of blood-stage malaria promotes Plasmodium-specific germinal center responses. JCI Insight. 3(2018), doi:10.1172/jci.insight.94142.

48. C. C. Kim, A. M. Baccarella, A. Bayat, M. Pepper, M. F. Fontana, FCRL5+ Memory B Cells Exhibit Robust Recall Responses. Cell Rep. 27, 1446–1460.e4 (2019).

49. M. Plantinga, M. Guilliams, M. Vanheerswynghels, K. Deswarte, F. Branco-Madeira, W. Toussaint, L. Vanhoutte, K. Neyt, N. Killeen, B. Malissen, H. Hammad, B. N. Lambrecht, Conventional and monocyte-derived CD11b(+) dendritic cells initiate and maintain T helper 2 cell-mediated immunity to house dust mite allergen. Immunity. 38, 322–335 (2013).

50. R. Gieseler, D. Heise, A. Soruri, P. Schwartz, J. H. Peters, In-vitro differentiation of mature dendritic cells from human blood monocytes. Dev Immunol. 6, 25–39 (1998).

51. A. H. Achtman, M. Khan, I. C. M. MacLennan, J. Langhorne, Plasmodium chabaudi chabaudi infection in mice induces strong B cell responses and striking but temporary changes in splenic cell distribution. J. Immunol. 171, 317–324 (2003).

52. M. M. Stevenson, G. Kraal, Histological changes in the spleen and liver of C57BL/6 and A/J mice during Plasmodium chabaudi AS infection. Exp. Mol. Pathol. 51, 80–95 (1989).

53. L. Beattie, C. R. Engwerda, M. Wykes, M. F. Good, CD8+ T lymphocyte-mediated loss of marginal metallophilic macrophages following infection with Plasmodium chabaudi chabaudi AS. J. Immunol. 177, 2518–2526 (2006).

54. B. C. Urban, T. T. Hien, N. P. Day, N. H. Phu, R. Roberts, E. Pongponratn, M. Jones, N. T. H. Mai, D. Bethell, G. D. H. Turner, D. Ferguson, N. J. White, D. J. Roberts, Fatal Plasmodium falciparum malaria causes specific patterns of splenic architectural disorganization. Infect Immun. 73, 1986–1994 (2005).

55. E. T. Cadman, A. Y. Abdallah, C. Voisine, A.-M. Sponaas, P. Corran, T. Lamb, D. Brown, F. Ndungu, J. Langhorne, Alterations of splenic architecture in malaria are induced independently of Toll-like receptors 2, 4, and 9 or MyD88 and may affect antibody affinity. Infect. Immun. 76, 3924–3931 (2008).

56. G. J. Keitany, K. S. Kim, A. T. Krishnamurty, B. D. Hondowicz, W. O. Hahn, N. Dambrauskas, D. N. Sather, A. M. Vaughan, S. H. I. Kappe, M. Pepper, Blood Stage Malaria Disrupts Humoral Immunity to the Pre-erythrocytic Stage Circumsporozoite Protein. Cell Rep. 17, 3193–3205 (2016).

57. V. Ryg-Cornejo, L. J. Ioannidis, A. Ly, C. Y. Chiu, J. Tellier, D. L. Hill, S. P. Preston, M. Pellegrini, D. Yu, S. L. Nutt, A. Kallies, D. S. Hansen, Severe Malaria Infections Impair Germinal Center Responses by Inhibiting T Follicular Helper Cell Differentiation. Cell Rep. 14, 68–81 (2016).

58. J. Rivera-Correa, M. F. Yasnot-Acosta, N. C. Tovar, M. C. Velasco-Pareja, A. Easton, A. Rodriguez, Atypical memory B-cells and autoantibodies correlate with anemia during Plasmodium vivax complicated infections. PLoS Negl Trop Dis. 14, e0008466 (2020).

59. J. Rivera-Correa, M. S. Mackroth, T. Jacobs, J. Schulze Zur Wiesch, T. Rolling, A. Rodriguez, Atypical memory B-cells are associated with Plasmodium falciparum anemia through anti-phosphatidylserine antibodies. Elife. 8, e48309 (2019).

60. C. Fernandez-Arias, J. Rivera-Correa, J. Gallego-Delgado, R. Rudlaff, C. Fernandez, C. Roussel, A. Götz, S. Gonzalez, A. Mohanty, S. Mohanty, S. Wassmer, P. Buffet, P. A. Ndour, A. Rodriguez, Anti-Self Phosphatidylserine Antibodies Recognize Uninfected Erythrocytes Promoting Malarial Anemia. Cell Host Microbe. 19, 194–203 (2016).

61. L. R. Sardinha, M. R. D’Império Lima, J. M. Alvarez, Influence of the polyclonal activation induced by Plasmodium chabaudi on ongoing OVA-specific B- and T-cell responses. Scand J Immunol. 56, 408–416 (2002).

62. C. D. Thouvenel, M. F. Fontana, J. Netland, A. T. Krishnamurty, K. K. Takehara, Y. Chen, S. Singh, K. Miura, G. J. Keitany, E. M. Lynch, S. Portugal, M. C. Miranda, N. P. King, J. M. Kollman, P. D. Crompton, C. A. Long, M. Pancera, D. J. Rawlings, M. Pepper, Multimeric antibodies from antigen-specific human IgM+ memory B cells restrict Plasmodium parasites. J Exp Med. 218, e20200942 (2021).

63. M. J. Boyle, L. Reiling, G. Feng, C. Langer, F. H. Osier, H. Aspeling-Jones, Y. S. Cheng, J. Stubbs, K. K. A. Tetteh, D. J. Conway, J. S. McCarthy, I. Muller, K. Marsh, R. F. Anders, J. G. Beeson, Human Antibodies Fix Complement to Inhibit Plasmodium falciparum Invasion of Erythrocytes and Are Associated with Protection against Malaria. Immunity. 42, 580–590 (2015).

64. D. Donati, L. P. Zhang, Q. Chen, A. Chêne, K. Flick, M. Nyström, M. Wahlgren, M. T. Bejarano, Identification of a Polyclonal B-Cell Activator in Plasmodium falciparum. Infect Immun. 72, 5412–5418 (2004).

65. A. Baccarella, M. F. Fontana, E. Chen, C. C. Kim, Toll-like receptor 7 mediates early innate immune responses to malaria. Infect. Immun. (2013), doi:10.1128/IAI.00923-13.

66. S. Sharma, R. B. DeOliveira, P. Kalantari, P. Parroche, N. Goutagny, Z. Jiang, J. Chan, D. C. Bartholomeu, F. Lauw, J. P. Hall, G. N. Barber, R. T. Gazzinelli, K. A. Fitzgerald, D. T. Golenbock, Innate immune recognition of an AT-rich stem-loop DNA motif in the Plasmodium falciparum genome. Immunity. 35, 194–207 (2011).

67. X. Wu, N. M. Gowda, S. Kumar, D. C. Gowda, Protein-DNA complex is the exclusive malaria parasite component that activates dendritic cells and triggers innate immune responses. J. Immunol. 184, 4338–4348 (2010).

68. J. Rivera-Correa, J. J. Guthmiller, R. Vijay, C. Fernandez-Arias, M. A. Pardo-Ruge, S. Gonzalez, N. S. Butler, A. Rodriguez, Plasmodium DNA-mediated TLR9 activation of T-bet+ B cells contributes to autoimmune anaemia during malaria. Nat Commun. 8, 1282 (2017).

69. J. Wipasa, C. Suphavilai, L. C. Okell, J. Cook, P. H. Corran, K. Thaikla, W. Liewsaree, E. M. Riley, J. C. R. Hafalla, Long-Lived Antibody and B Cell Memory Responses to the Human Malaria Parasites, Plasmodium falciparum and Plasmodium vivax. PLOS Pathogens. 6, e1000770 (2010).

70. P. Jahnmatz, D. Nyabundi, C. Sundling, L. Widman, J. Mwacharo, J. Musyoki, E. Otieno, N. Ahlborg, P. Bejon, F. M. Ndungu, A. Färnert, Plasmodium falciparum-Specific Memory B-Cell and Antibody Responses Are Associated With Immunity in Children Living in an Endemic Area of Kenya. Front Immunol. 13, 799306 (2022).

71. D. Chandramohan, I. Zongo, I. Sagara, M. Cairns, R.-S. Yerbanga, M. Diarra, F. Nikièma, A. Tapily, F. Sompougdou, D. Issiaka, C. Zoungrana, K. Sanogo, A. Haro, M. Kaya, A.-A. Sienou, S. Traore, A. Mahamar, I. Thera, K. Diarra, A. Dolo, I. Kuepfer, P. Snell, P. Milligan, C. Ockenhouse, O. Ofori-Anyinam, H. Tinto, A. Djimde, J.-B. Ouédraogo, A. Dicko, B. Greenwood, Seasonal Malaria Vaccination with or without Seasonal Malaria Chemoprevention. N Engl J Med. 385, 1005–1017 (2021).

72. B. W. Huang, E. Pearman, C. C. Kim, Mouse Models of Uncomplicated and Fatal Malaria. Bio Protoc. 5(2015).

73. J. J. Taylor, K. A. Pape, M. K. Jenkins, A germinal center-independent pathway generates unswitched memory B cells early in the primary response. J. Exp. Med. 209, 597–606 (2012).

